# Similar processing of novelty in rat dorsal and intermediate CA1 despite differences in spatial tuning

**DOI:** 10.1101/2025.11.03.686060

**Authors:** Shang Lin Tommy Lee, Ryan Troha, Jinah Yoon, Aditi Anam, Kori-Anne Citrin, Qingli Hu, Miriam Katz, Mitchel Kuperstein, Divya Subramanian, David Katz, Kavya Katugam, Megan Pattoli, Stephanie Vu, Ian H. Stevenson, Etan J. Markus

## Abstract

Elucidating the fundamental neural mechanisms of hippocampal information processing is necessary for understanding memory formation and related brain disorders. Differences in hippocampal genetics, anatomy, and connectivity across the longitudinal axis suggest functional heterogeneity in this structure. The dorsal pole is suggested to be primarily involved in spatial processing, whereas the ventral pole is implicated in emotional processing. Connectivity and genetic studies show this functional segregation is more prominent near their respective poles and weaker toward the intermediate region. The current study compares the firing properties of CA1 cells in the dorsal and intermediate regions of the hippocampus during spatial or social/odor novelty. We measured basic spatial properties, firing rate response, and remapping to the spatial re-configuration of a linear track or the social/odor presentation of a novel male conspecific, female bedding, or coyote urine. Behaviorally, the average rat exhibited slower maze running latencies during the novel spatial manipulation and spent more time adjacent to the chamber containing the novel social/odor stimulus. As previously shown, dorsal cells had fewer, smaller place fields, and higher spatial information content than intermediate cells in the hippocampus. Despite the differences in place field characteristics, cells in both regions responded similarly to spatial and social/odor manipulations. Taken together, these data support the differentiation of some functions, together with an overlap of other functions progressing along the longitudinal axis, which may facilitate the integration of information throughout the hippocampus.

## Introduction

The link between episodic memory formation and the hippocampus is well established. Far less is known regarding the segregation and integration of processing among different regions of the hippocampus. Differences in hippocampal genetics, anatomy, and connectivity across the longitudinal axis suggest functional heterogeneity in the rodent dorsal (dHP), intermediate (iHP), and ventral (vHP) hippocampus (Strange et al., 2014). The dorsal pole is suggested to be primarily involved in spatial processing, whereas, the ventral pole is implicated in emotional processing (Fanselow & Dong, 2010). The anatomical and connectivity differences seem most prominent near their respective poles and weaken toward the iHP. Thus, the degree to which the iHP processes information similar to the dHP warrants further investigations.

In rodents, traditional views of the dorsal-ventral dichotomy are based on observations of differential inputs to the entorhinal cortex, which projects topographically to distinct regions of the hippocampal long axis (Burwell & Amaral, 1998; Dolorfo & Amaral, 1998; Suzuki & Amaral, 1994; Witter & Groenewegen, 1984). Outputs of dHP lead to the dorsal subiculum, retrosplenial cortex, dorsal lateral septum, and mammillary body, whereas, outputs of vHP lead to the olfactory, ventral lateral septum, amygdala, medial prefrontal cortex, and subcortical structures associated with the hypothalamic-pituitary-adrenal axis (Chiba, 2000; Gergues et al., 2020; Pitkänen et al., 2000; Risold & Swanson, 1996; Van Groen & Wyss, 1990). Connectivity to NAc follows a gradient of dHP to core and vHP to shell regions (Groenewegen et al., 1987). Unlike dHP, iHP and vHP does not directly project to retrosplenial cortex (Cenquizca & Swanson, 2007). Furthermore, there is a gradient of connectivity from iHP to prelimbic and vHP to infralimbic mPFC (Jay & Witter, 1991). Unlike the vHP, iHP and dHP lack direct projections to the amygdala and hypothalamus (Kishi et al., 2006). Moreover, vHP projects to the medial olfactory nucleus, whereas iHP projects to the lateral anterior olfactory nucleus (Aqrabawi & Kim, 2018). Taken together, these results suggest a degree of functional segregation along the hippocampal long axis.

Dorsal hippocampus damage results in impaired spatial processing (Holt & Maren, 1999; E. Moser et al., 1993; Pothuizen et al., 2004), whereas, ventral regions are necessary for modulating anxiety and odor-related behavioral responses (Bannerman et al., 2004; Bast et al., 2001; Maren & Holt, 2004; Pentkowski et al., 2006; Richmond et al., 1999; Weeden et al., 2015). Dorsal hippocampus neurons have small precise location specificity, whereas, intermediate to ventral neurons exhibited broader place fields, decreased spatial information content, and lower mean firing rates (Jung et al., 1994; Kjelstrup et al., 2008; Komorowski et al., 2013; Maurer et al., 2005; O’Keefe & Dostrovsky, 1971; Poucet et al., 1994; Royer et al., 2010). Conversely, ventral units show selective firing during social, olfactory, and fearful encounters (Ciocchi et al., 2015; Jimenez et al., 2018; Mikulovic et al., 2018; Rao et al., 2019; M. E. Wang et al., 2012). Importantly, hippocampal associational projections from CA3 pyramidal cells and dentate hilar mossy cells project to approximately two-thirds of the dorsoventral extent of the hippocampus, suggesting a degree of functional integration between dorsal, intermediate, and ventral regions (Amaral & Witter, 1989; Fricke & Cowan, 1978). Hence, intermediate hippocampus may be critical for integrating visuospatial information in dorsal hippocampus with behavioral control mechanisms in ventral regions (Bast et al., 2009). In support of functional integration, we previously showed dorsal and intermediate/ventral hippocampus to be interdependent for precise spatial navigation (Lee et al., 2019). Further, both dorsal or intermediate/ventral hippocampus disruption both eliminated a prospective bias during a sequential order spatial memory task (Lee et al., 2023). Notably, in humans, there is also evidence for different types of processing in the iHP (Dalton et al., 2018; Evensmoen et al., 2015; Poppenk et al., 2013). The current study compared and contrasted unit activity in dorsal CA1 (dCA1) and intermediate CA1 (iCA1). Specifically, we focused on the following issues:

1. Firing properties of dCA1 and iCA1 cells under stable (familiar) conditions.
2. Changes in firing properties of dCA1 and iCA1 cells in response to a novel event in the environment.

The current study shows both brain areas responded similarly to novelty despite differences in their spatial tuning, suggesting they are sending out similar information but to distinct downstream targets.

## Materials and Methods

### Subjects

Twenty-two male Fischer (F-344) rats (Envigo), arriving at four months of age, were housed in a colony maintained at 23 °C with 12-h light/dark cycle (lights on at 8:00 am). Rats were fed standard rat chow (Teklad Global Rodent Diets) in addition to chocolate sprinkles (UConn Dairy Bar) and Dustless Precision Pellets (F0021; Bio-Serv) for reward during behavioral performance. All rats were single-housed, food restricted to 85% of their free feeding weight, and had *ad libitum* access to water. All experiments were conducted in accordance with University of Connecticut Institutional Animal Care and Use Committee.

### Hyperdrive construction

Hyperdrive recording devices contained 16 independently moveable tetrodes. Each tetrode comprised of four polyamide-coated 14-μm nickel-chrome wires (Sandvik) twisted together. Each tetrode was inserted into polymicro tubing (Polymicro Technologies), which was nested within a 30-gauge stainless steel cannula. One full turn of each independently moveable tetrode carrier screw was approximately 318 μm. Electrodes were gold plated to reduce single channel impedances to approximately 150-500 kΩ at 1 kHz by passing current through the wires while the electrode tips were immersed in a mixture of gold plating (PAS-5355; Sifco) and polyethylene glycol solution (PEG 8000 MW; Sigma-Aldrich) (Ferguson et al., 2009).

### Pre-surgery behavioral training

Rats were trained to run on a linear track maze (130 cm x 10 cm) composed of two arms, one stationary arm and one rotatable arm. Rats ran back and forth to both ends of the linear track to receive Bio-Serv pellet food rewards from automated feeders (Fig. 1). Rats were trained for three 10 min sessions per day with five min inter-session-intervals. Rats were trained until they reach a criterion of at least 20 trials (10 back and forth) in each of the three sessions per day on at least two consecutive days of training. After reaching criterion, rats received the surgical hyperdrive implantation.

**Figure 1.**
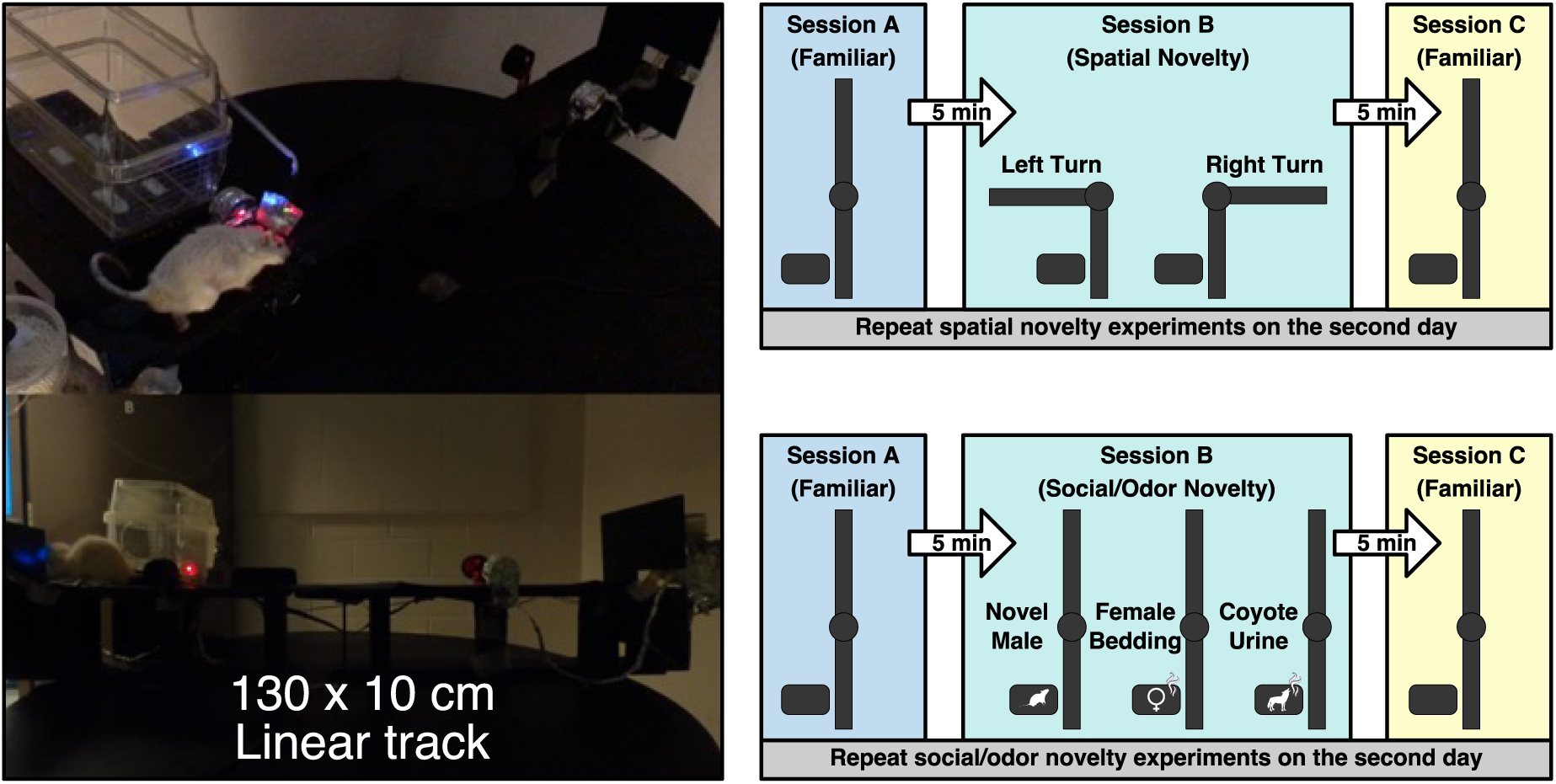
Behavioral paradigm. Photos of a hyperdrive implanted rat running on a 130 x 10 cm linear track. During novelty days, rats ran on the maze across three sessions (Session A “Familiar” → Session B “Novelty” → Session C “Familiar”). Each session was preceded and followed by five-minute rest periods within a cage next to the maze. Novelty events included spatial trajectory (left or right turn) or social/odor (novel male conspecific, female bedding, or coyote urine) manipulations. The same manipulation was repeated the following day (Day 2, Session B).

### Surgery

Male F-344 rats, approximately 10 months of age, were implanted with tetrode hyperdrives using sterile surgical techniques. Antibiotics Enrofloxacin (10 mg/kg, s.c.) was injected two hours prior to the first incision, every two hours during surgery, and post-operatively twice per day for 48 hours. At the time of surgery, rats were anesthetized with isoflurane (3% in O_2_ during induction, 1-3% in O_2_ during maintenance). After confirmation of anesthesia, meloxicam (Metacam, 1 mg/kg, s.c.) was injected, each rat was placed in a stereotaxic apparatus with blunt ear bars, ophthalmic ointment was applied on their eyes, and their scalp was shaved and wiped with 2% chlorhexidine, 70% ethanol, and Betadine. A small midline scalp incision was made, and a craniotomy drilled over the right hippocampus using rat brain atlas (Paxinos & Watson, 2007) coordinates -4.7 mm anteroposterior and 3.0 mm mediolateral from bregma, as well as -6.8 mm anteroposterior and 5.8 mm mediolateral from bregma, to allow an array of electrode to target both the dorsal and intermediate hippocampus simultaneously. Five stainless steel screws were secured to the skull. The initial length of the tetrodes targeting dorsal hippocampus extended 1 mm beyond the hyperdrive base and tetrodes targeting ventral hippocampus extended 5 mm beyond the hyperdrive base. After the dura was carefully removed, the hyperdrive was positioned above the craniotomy, and the extended tetrodes penetrated the brain surface until the hyperdrive base reached the brain surface. Silicone elastomer (Kwik-sil; WPI) covered the remaining exposed brain within the craniotomy. Dental acrylic (B1334; Ortho-Jet BCA) was applied to the hyperdrive base and screws to anchor the device. After hyperdrive implantation, two additional injections of Meloxicam were administered post-operatively at 24 and 48 h after surgery. Rats received ten days of post-operative recovery before being food-restricted to 85% of their free feeding weight and behavioral testing.

### Post-surgery behavioral re-training

Rats were re-trained to run on the linear track to receive Bio-Serv pellet food rewards from two automated feeders at the ends of the maze. Before each day’s training session, rats were placed in a small transparent cage (rest box; 27.3 cm x 16.5 cm x 21.6 cm) with an open top and an 8 cm diameter circular opening (covered by a metal wire mesh) facing the maze apparatus. After five min in the maze apparatus (pre-maze rest), rats were re-trained for three 10 min maze sessions per day with five min inter-session-intervals inside the rest box. Upon the completion of the third session, rats were placed in the rest box for another five min (post-maze rest). Rats were re-trained on the linear track maze until they reach a criterion of at least 20 trials (10 back and forth) in each of the three sessions per day on at least two consecutive days of training. Tetrodes were gradually lowered until hippocampal single units and characteristic LFP patterns (sharp-wave ripples; theta rhythm) can be identified. Novelty experiments began when rats reach criteria and single-units were well isolated.

### Novelty experiment conditions

Spatial novelty included spatial re-configuration of the linear track to a 90 left or right turn. Social/odor novelty included presentation of a novel male rat, female bedding, or coyote urine in a small cage next to the linear track.

The novel rat was a male conspecific placed in the small cage with the opening adjacent to the linear track and at a distance that prevented the rats from touching vibrissa. The female bedding experiment used a mix of novel female conspecific rat bedding from multiple cages. Coyote urine (Bare Ground, Framingham, MA) was prepared by placing three drops on top of cotton balls within a hexagonal polystyrene tray. Running latency during novelty days was measured as the animal’s roundtrip journey (excluding time spent eating the food reward). For the social/odor manipulations, an experimenter monitored the animals and noted when the rat first noticed (e.g. scanning, pausing, rearing) the novel stimulus. The behavioral flag was done blind to the animal’s neuronal activity. All novelty conditions were presented across two consecutive days (Day 1 and 2). Animals could belong to more than one novelty experiment conditions. The ordering of the novelty experiment conditions was consistent across all animals in the following sequence: left turn, right turn, novel male conspecific, female bedding, and coyote urine.

### Data acquisition

The electrode interface board (EIB-72-QC-Large; Neuralynx) on the hyperdrive was connected to a multichannel, unity gain headstage (HS-72-QC-LED; Neuralynx). The output of the headstage was conducted via two lightweight tether cables to a digital data acquisition system that processed the signals (Digital Lynx SX; Neuralynx). Continuously sampled raw broadband data (between 0.1 and 8,000 Hz) were acquired and saved at 32 kHz prior to being processed. The headstage had an array of light-emitting diodes (three blue and three red LEDs) to track the rat’s position and head direction. XY position coordinates of both diode arrays were sampled at 29.97 Hz using an overhead video tracking system (Neuralynx). Speed was calculated by taking the finite difference between successive position coordinates followed by a low-pass filter (cutoff = 0.25 Hz) to minimize movements and other movement related artifacts.

### Spike sorting

MountainSort4 automated spike sorting algorithm was used to bandpass filter (600-6,000 Hz) the raw data, detect spikes, and isolate single-units (Chung et al., 2017). Cluster quality thresholds were applied to only include single units with greater than 95% isolation, less than 3% noise overlap, and greater than 1.5 peak signal-to-noise ratio (SNR). The isolation metric quantifies how well separated the cluster is from other nearby clusters, the noise overlap metric estimates the fraction of above-threshold events not associated with true firings in a clustered unit, and the peak SNR is defined as the peak absolute amplitude of the average waveform divided by the peak standard deviation. This was followed by manual curation of clusters based on visual inspection of waveforms, auto-correlograms, and cross-correlograms to obtain well-isolated single units. Only data from stable recordings were included in the analysis. Stability of a cluster was defined as the average spike amplitude of the last session being within ± one standard deviation from the average and distribution of spike amplitudes of the first session.

### Histology

At the conclusion of the experiments, rats were euthanized with CO_2_, and intracardially perfused with phosphate buffered saline (PBS) followed by 4% paraformaldehyde in PBS. Brains were post-fixed for at least 24 h in 4% paraformaldehyde in PBS. Brains were sectioned 40-75 μm thick with a vibratome (VT-1000S; Leica), mounted on glass microscope slides, stained with thionin, and covered with DPX slide mounting medium. Tetrode tracks were examined using a light microscope to determine locations of recorded single-units.

### Proportional firing rate

Mean firing rate (total number of spikes divided by total duration of maze session) was computed for each single unit during the maze sessions. The proportional firing rate was calculated as the mean firing rate in a single maze session divided by the mean firing rate across the three maze sessions.

### Classification of putative cell type

Identification of putative interneurons were based on single-units with spike width <350 μs and firing rate >7.5 Hz, and were excluded from rate map analyses.

### Classification of dorsal and intermediate hippocampal units

The division between dCA1 and iCA1 was based on the criteria used by (Jin & Lee, 2021). Dorsal CA1 units were defined by tetrode tips found anterior to -4.2 mm anteroposterior from bregma using the rat brain atlas (Paxinos & Watson, 2007). Intermediate CA1 units were defined by tetrode tips posterior to -4.2 mm anteroposterior and above 5.5 mm dorsoventral from bregma (Fig. 2).

**Figure 2.**
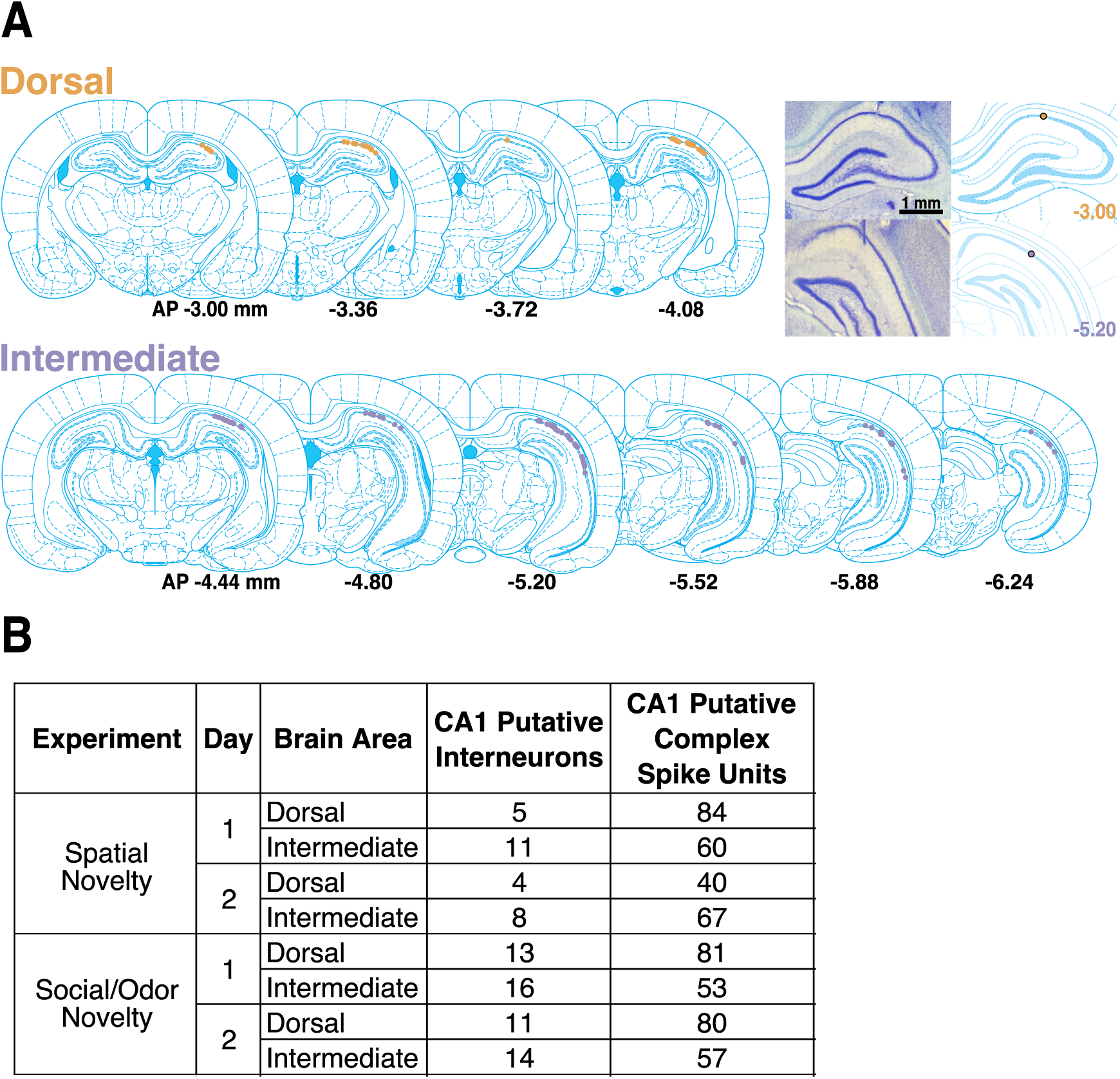
Histology summary of CA1 tetrode locations and cutoff criteria. **(A)** Histological reconstruction of dorsal and intermediate CA1 units overlayed on the rat brain atlas (Paxinos & Watson, 2007). Dorsal units were defined by tetrode tips found anterior to -4.2 mm anteroposterior (AP) from bregma. Intermediate units were defined by tetrode tips posterior to -4.2 mm anteroposterior and above 5.5 mm dorsoventral from bregma. Example Nissl-stained histological verification of tetrode placement within the dorsal and intermediate hippocampus are shown on the right. **(B)** Table with the number of recorded CA1 putative interneurons or CA1 putative complex spike units across experiments, days, and brain areas.

### Firing rate maps and place field characteristics

To analyze place selectivity, position data was divided into 0.3 x 0.3 cm bins. The total number of spikes that occur in a given spatial location bin was summed and smoothed with a 20 x 20 bin Gaussian filter with a standard deviation of approximately 2.5 bins. The total amount of time that the rat spent in a given spatial location bin was also summed and smoothed using the same parameters. Smoothed rate maps were calculated as the smoothed total number of spikes divided by the smoothed time spent in each spatial location bin. Bins with insufficient sampling (<3 visits or <1 ms in time spent) were regarded as unvisited and were not included in the rate map. Rate maps were only calculated for periods when the rat is moving at a robust speed (> 6 cm/s) to control for possible influence of stationary periods. Analyses of rate maps were separated into two directions to control for place cell directional selectivity on linear track mazes (Markus et al., 1995). Place fields were defined as an area of at least 200 contiguous bins with a firing rate per bin >20% of the peak firing rate of the cell during the maze session. Proportion of active bins was calculated as the number of bins >20% of the peak firing rate of the cell during the maze session divided by the total number of bins in the maze session.

### Spatial correlations

Place field stability was calculated by measuring the correlation between the firing rate maps of a cell within and across maze sessions on a bin-by-bin basis. For spatial novelty, the firing rate map of the re-configured maze arm was rotated clockwise or counter-clockwise by 90 degrees prior to correlation analyses so they can be compared to linear firing rate maps on a bin-by-bin basis. The prerequisite for performing a rate map correlation was a minimum of one place field during at least one of the maze sessions of a recording. Correlations between maze sessions were computed only if the cell fired at least 20 spikes in each maze session. Within-maze session correlations were calculated by comparing the firing rate maps of the first five trials and the last five trials of the maze session. Correlations within maze sessions were computed only if the cell fired at least 10 spikes in each of the five trials.

### Spatial information content

The spatial information (bits) per spike for each neuron was calculated using the smoothed rate maps, in terms of the spatial information a single spike conveys about the rat’s location. Spatial information content of spike discharge was calculated using the following formula:

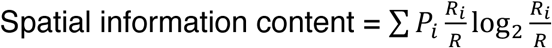

where *i* is the bin number, *P_i_* is the probability of occupancy in bin *i* (total time spent at bin *i* divided by total time spent through the maze session), *R_i_* is the mean firing rate for bin *i*, and *R* is the mean firing rate of the cell during the maze session (Skaggs et al., 1993).

### Relative place field characteristics

The change in basic place field characteristics was calculated as the value for “Maze B to A” = Session B/(Session B + Session A), “Maze B to C” = Session B/(Session B + Session C), and “Maze C to A” = Session C/(Session C + Session A). This was done for the total number of place fields, largest place field’s size (pixels), percent of pixels above 20% of the peak firing rate, and average firing rate within the largest place field.

## Results

Our experiment trained male rats to retrieve food rewards on both ends of a familiar linear track maze (Fig. 1). Rats were implanted with tetrodes targeting the hippocampus (Fig. 2), and recorded across three behavioral sessions (A, B, and C). During sessions A and C, rats were in the familiar linear track, while in B they experienced a novel condition. Twenty rats with well-isolated single-units experienced a novel spatial maze left turn (rotated 90deg counterclockwise), eighteen rats experienced a novel spatial maze right turn (rotated 90deg clockwise), eighteen rats experienced a novel rat cue, seventeen rats experienced a novel female bedding odor cue, and seventeen rats experienced a novel coyote urine odor cue. To examine the differences between the first exposure to a novel situation (Session B) and subsequent exposures, rats were recorded across two sequential days with the same pattern of three sessions. Each session ended when the rat completed 10 roundtrips on the maze. Before and after each session, rats spent five minutes in a rest box next to the maze.

We compared and contrasted unit activity in dCA1 and iCA1 as rats were exposed to spatial or social/odor novelty. To exclude putative interneurons, units were sorted by calculating spike width, average firing rate, and spatial information content (Fig. 3). As with previous studies (Markus et al., 1994), we found that units with narrow spike waveforms and high firing rates (putative interneurons) tend to have lower spatial information content, while putative principal cells with broad spike waveforms are often place selective.

**Figure 3.**
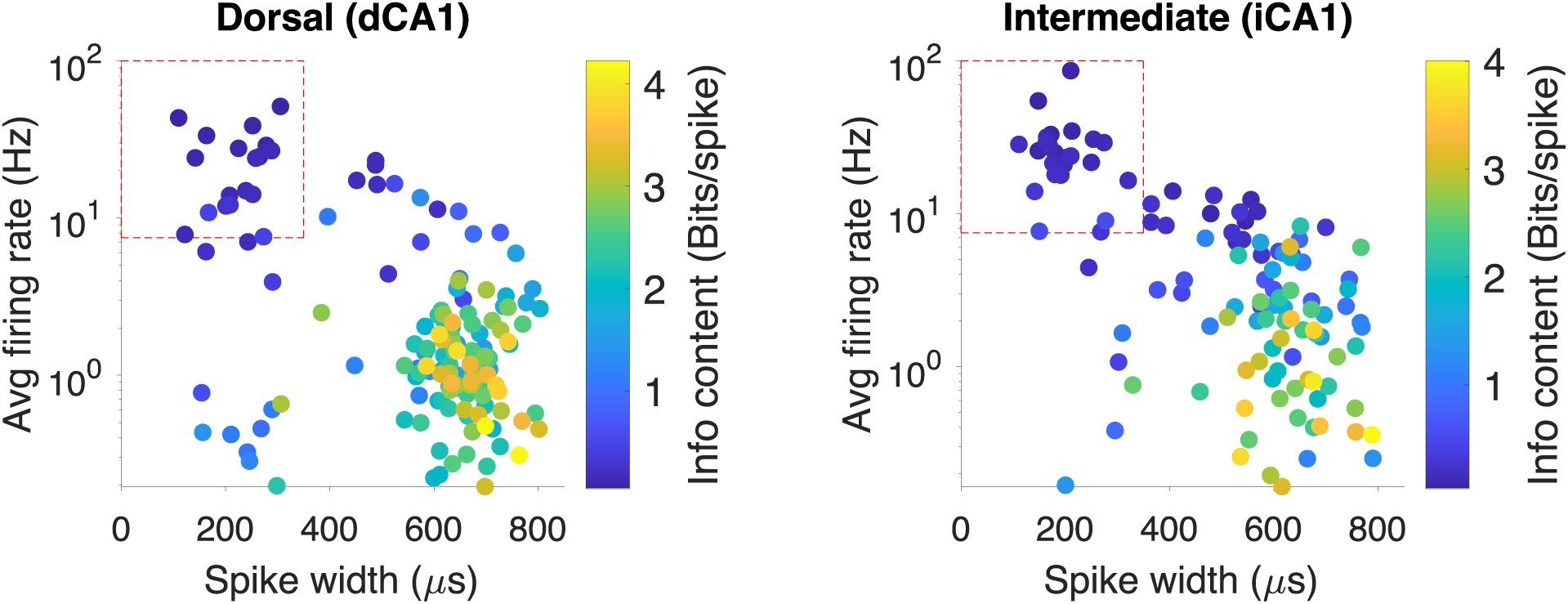
Putative CA1 interneurons and complex spike units. Dorsal and intermediate CA1 putative interneuron and complex spike units are classified in a scatter plot of spike width x firing rate x spatial information content. Identification of putative interneurons were based on single-units with spike width <350 μs and firing rate >7.5 Hz, and were excluded from rate map analyses.

Raw place field characteristics of rate maps were compared between dCA1 and iCA1 place cells across the first day of all spatial and social/odor novel conditions (Fig. 4). Independent samples t-tests between dCA1 and iCA1 found place cells in dCA1 exhibited higher spatial information content, fewer total number of place fields, smaller place field size, and lower percentage of active pixels above 20% of the peak firing rate (*t*_202_ = 4.05, *p* < 0.001; *t*_202_ = -2.67, *p* < 0.01; *t*_202_ = -3.36, *p* < 0.001; *t*_202_ = -2.87, *p* < 0.01, respectively).

**Figure 4.**
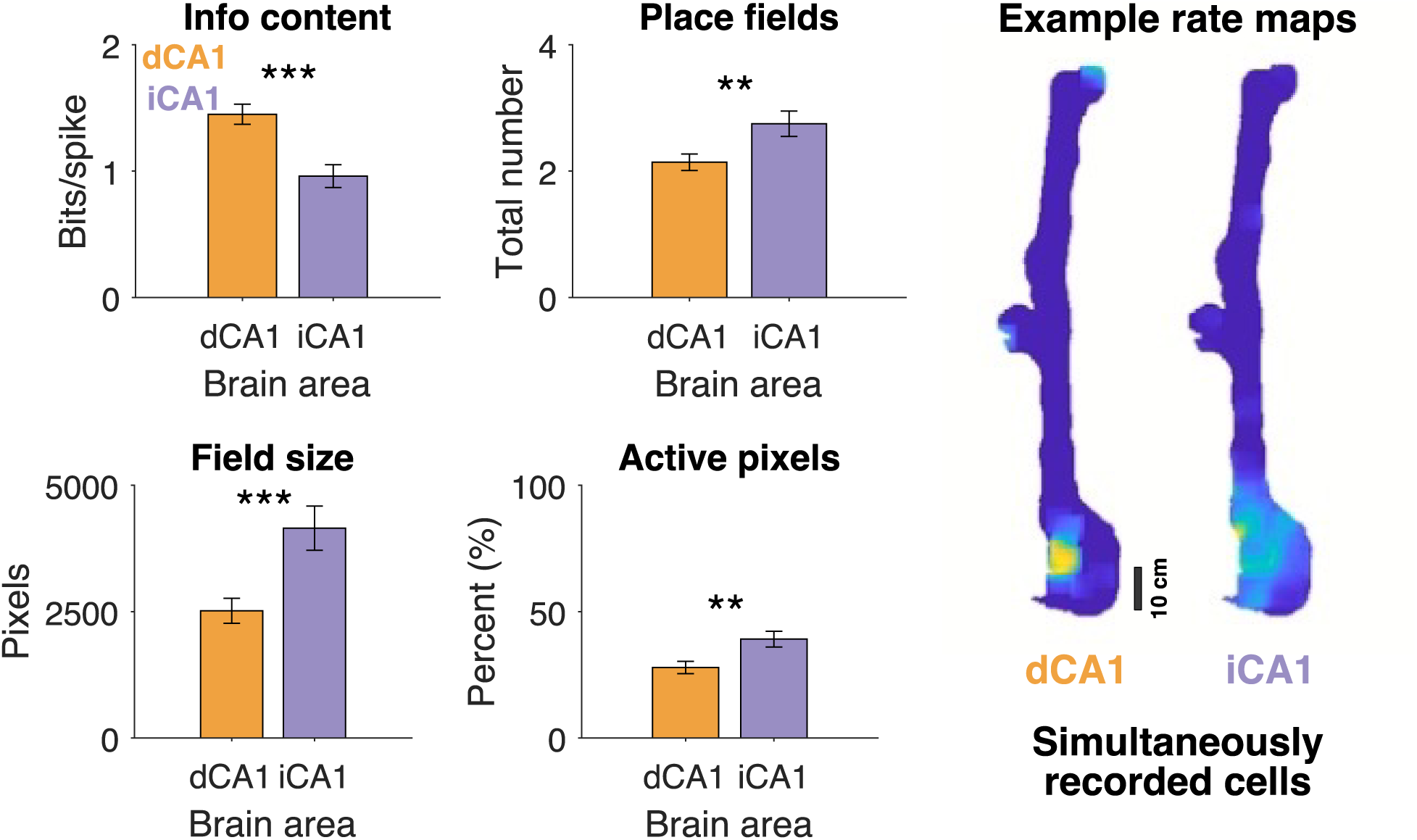
Raw place field characteristics of dorsal and intermediate CA1 cells. Raw place field characteristics were computed for stable cells during Session A (“Familiar”). The following characteristics were calculated: Spatial information content, total number of place fields on the maze, largest place field’s size (pixels), and percent of pixels above 20% of the peak firing rate. Error bars denote standard error of the mean. Example place fields from two simultaneously recorded dCA1 and iCA1 cells are shown. Yellow color indicates the peak (higher) firing rate.

Our results, similar to previous findings, suggest dCA1 place cells have smaller, precise, place fields with more spatial information content than iCA1 place cells. Further, these results suggest iCA1 cells were more active in multiple areas on the maze.

The number of units recorded for each experiment, day, and brain area are presented (Fig. 2B). We found no systematic differences between left/right turn experiments and no systematic difference between novel rat/female bedding/coyote urine experiments. For further analyses we thus combined experiments into two general conditions: “Spatial Novelty” and “Social/Odor Novelty.”

### Running latency under familiar and novel conditions – Spatial novelty

We found that the latency across sessions A, B, and C differed substantially for the spatial novelty experiments (Fig. 5A left; repeated-measures two-way ANOVA, *F*_2, 216_ = 20.37, *p*< 0.001). Average latency was longer in B vs. A or C (*p* < 0.001, post hoc comparisons using the Tukey HSD test), and there was no statistically significant difference between A vs. C (*p* > 0.10). Day 1 had longer latencies than the second day (*F*_1, 216_ = 8.23, *p* <0.01). There was also a statistically significant interaction of day and session (*F*_2, 216_ = 9.59, p < 0.001) where the long latency in session B was particularly long on the first day.

**Figure 5.**
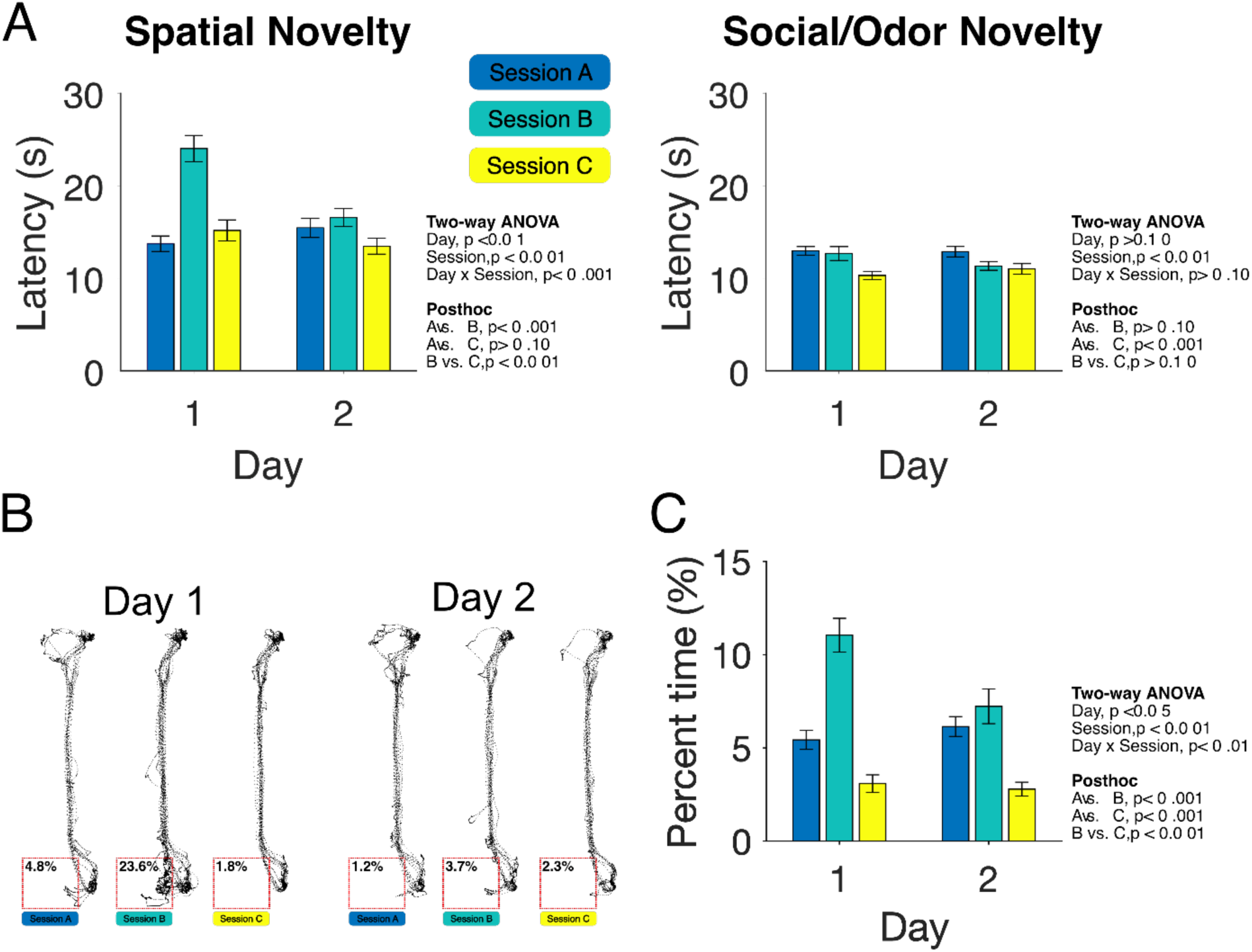
Running latency and trajectory under familiar and novel conditions. **(A)** Total roundtrip running latency during spatial (left panel) or social/odor (right panel) novelty manipulation days. Each novelty manipulation was repeated on a second day. Error bars denote standard error of the mean. **(B)** Example trajectories during a social/odor novelty experiment (Session B includes the presence of the social/odor cue). Red dotted line boxes represent the spatial zone directly adjacent to the social/odor cue. Percent time scores within each box represent the amount of time (%) the animal spent within the social/odor zone. **(C)** Average percent time animals spent within the social/odor zone across days and sessions. Error bars denote standard error of the mean.

### Running latency and trajectory under familiar and novel social/odor conditions

For the social/odor novelty conditions, we again found a statistically significant difference in running latency across sessions A, B, and C (Fig. 5A right; repeated-measures two-way ANOVA, *F*_2, 306_ = 8.45, *p* < 0.001), with the average latency was shorter on session C vs. A (*p* < 0.001, post hoc comparisons using the Tukey HSD test). There was no statistically significant difference between session B vs. session A or C (*p* > 0.10). The main effect for day and the interaction were not statistically significant (*p* > 0.10). As indicated in the Methods section, the experimenter noted if and when animals showed an overt response to the presence of the social/odor cue. Therefore, we examined the trajectory of the rats from that point. We found a statistically significant difference in the percent of time rats spent adjacent to the social/odor stimulus chamber across sessions for the social/odor novelty experiments (Fig. 5B and 5C; repeated-measures two-way ANOVA, *F*_2, 240_ = 44.65, *p* < 0.001). Rats spent a larger percent of their time adjacent to the chamber when it contained the social/odor stimulus (session B) vs. session A or C (*p* < 0.001 for both post hoc comparisons using the Tukey HSD test). Further, percent time spent adjacent to the stimulus chamber was lower in session C vs. A (*p* < 0.001). Day 1 had higher percent time spent adjacent to the stimulus chamber than Day 2 (*F*_1, 240_ = 4.46, *p* < 0.05). Lastly, there was a statistically significant interaction of day and session (*F*_2, 240_ = 6.56, *p* < 0.01) where the percent time spent adjacent to the social/odor stimulus in session B was higher on the first day. Taken together, these data show behavioral changes in response to the novel social/odor manipulation.

### Average proportional firing rate analysis – Spatial novelty

We recorded putative complex spike units in dCA1 (N=124) and in iCA1 (N=127) for animals (N=20) performing the spatial novelty experiment, and we recorded putative complex spike units in dCA1 (N=161) and in iCA1 (N=101) for animals (N=18) performing the social/odor novelty experiments. To compare changes in the average firing rate across days, sessions, and brain area we calculated the proportional firing rate for each unit by calculating the mean firing rate in a single maze session divided by the mean firing rate across the three maze sessions. A three-way ANOVA was conducted to examine the main effects and interactions of day, session, and brain area as they relate to the average proportional firing rate during spatial novelty (Fig. 6). There was a statistically significant difference in average proportional firing rate across sessions A, B, and C for the spatial novelty experiment (*F*_2, 672_ = 23.26, *p* < 0.001). Average proportional firing rate was higher in B vs. A (post hoc comparisons using the Tukey HSD; *p* < 0.001), lower in C vs. B (*p* < 0.001), and higher in C vs. A (*p* < 0.05). There was also a significant interaction of day and session, (*F*_2, 672_ = 4.39, *p* < 0.05). There was no statistically significant main effect of brain area, and all other main effects and interactions were not statistically significant (all *p* > 0.10).

**Figure 6.**
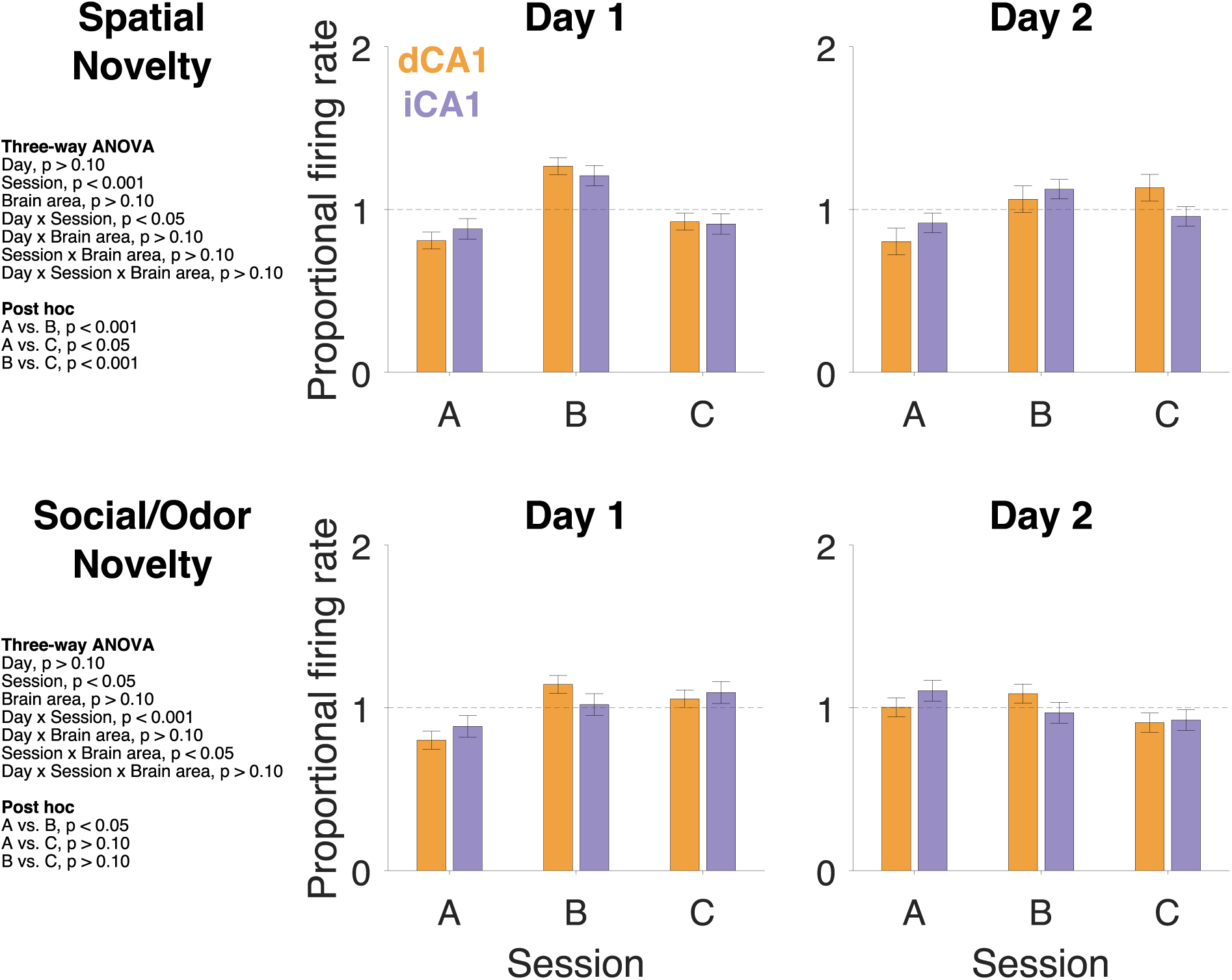
Proportional firing rate during novelty days. The proportional firing rate was calculated as the cell’s mean firing rate in a single maze session divided by the mean firing rate across the three maze sessions. Values > 1 indicate cell’s mean firing rate in a single maze session is higher than their mean firing rate across the three maze sessions. Error bars denote standard error of the mean.

### Average proportional firing rate analysis – Social/odor novelty

A three-way ANOVA was conducted to examine the main effects and interactions of day, session, and brain area as they relate to the average proportional firing rate during social/odor novelty (Fig. 6). There was a main effect for session (*F*_2, 549_ = 3.01, *p* < 0.05). Average proportional firing rate was higher in B vs. A (post hoc comparisons using the Tukey HSD; p < 0.05), and the other multiple comparisons across sessions were not statistically significant. There was a statistically significant two-way interaction of day and session (*F*_2, 549_ = 9.57, *p* < 0.001). There was also a statistically significant two-way interaction of session and brain area (*F*_2, 549_ = 3.21, *p* < 0.05). All other main effects and interactions were not statistically significant (all *p* > 0.10).

### Direct rate map correlations – Spatial novelty

Complex spike CA1 units, with a place field in at least one of the three sessions, were used in calculating direct rate map correlations (Fig. 7A). A three-way ANOVA was conducted to examine the main effects and interactions of day, sessions (A/B or A/C), and brain area as they relate to correlations across maze sessions (Fig. 7B). We found a statistically significant main effect for sessions, with correlations between A/C to higher than those for A/B for the spatial novelty task (*F*_1, 384_ = 8.91, *p* < 0.01). We additionally found that Day 2 had a higher average correlation across maze sessions than Day 1 (*F*_1,384_ = 7.30, *p* < 0.01). All other main effects (e.g. between dCA1 and iCA1) and interactions were not statistically significant (all *p* > 0.10).

**Figure 7.**
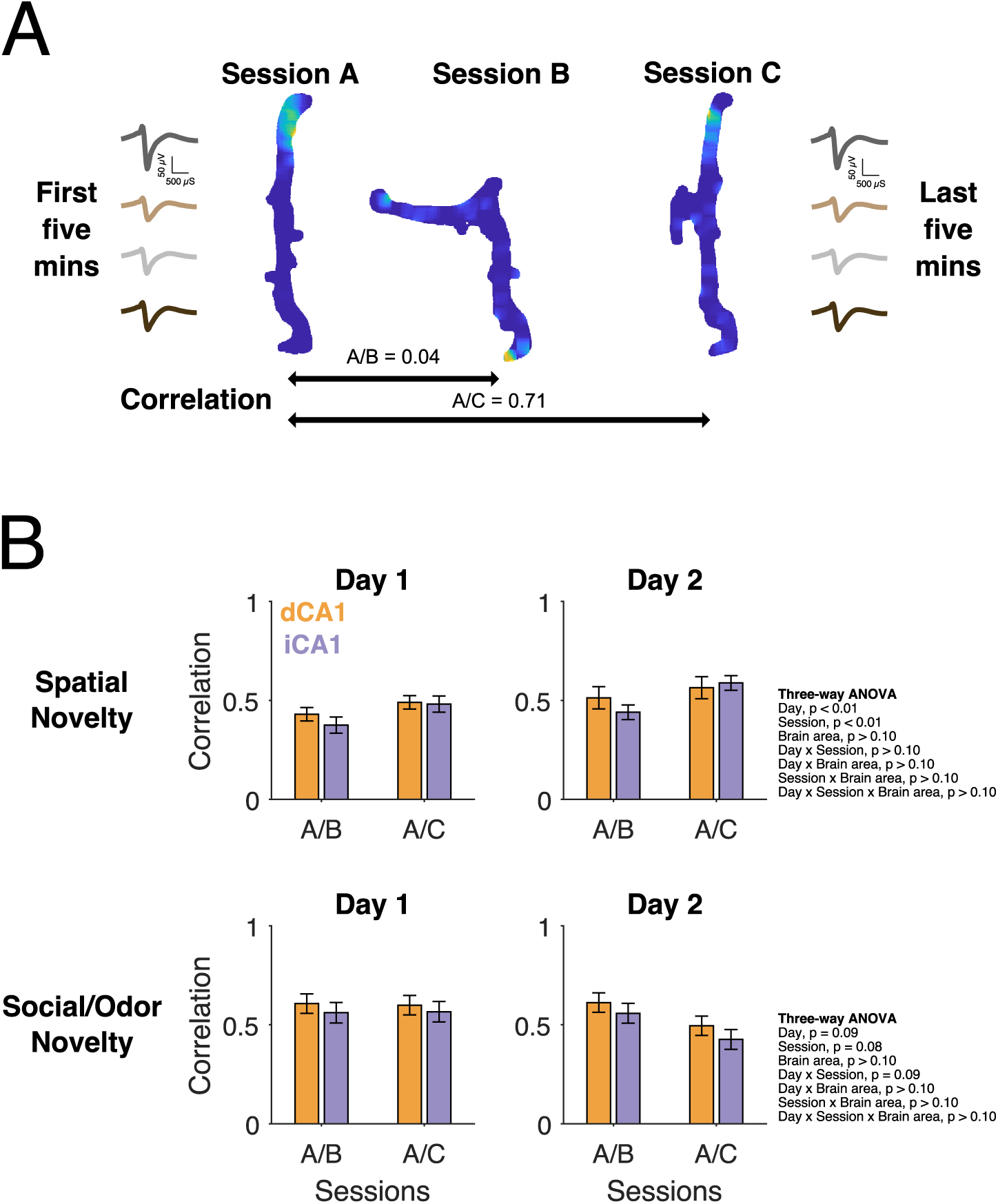
Rate map correlations across sessions. **(A)** Example of stable average waveforms from a dCA1 cell across the four channels of a tetrode are shown for the first and last five minutes of the recording day. Spatial firing rate maps from the cell across sessions and comparison of correlations are shown. Rate map correlation between linear rate maps in Sessions A and C are computed on a bin-by-bin basis. Rate map correlation between the linear rate map in Session A and the spatial novelty left turn rate map in Session B are computed, after rotating the re-configured left turn arm 90 degrees counterclockwise, on a bin-by-bin basis. Yellow color indicates the peak (higher) firing. **(B)** Average rate map correlations of spatial or social/odor novelty days are compared across days, sessions, and brain areas. Error bars denote standard error of the mean.

### Direct rate map correlations – Social/odor novelty

A three-way ANOVA was conducted to examine the main effects and interactions of day, sessions (A/B or A/C), and brain area as they relate to correlations across maze sessions (Fig. 7B). The main effects of day (*F*_1, 242_ = 2.94, *p* = 0.09) and session (*F*_1, 242_ = 3.17, *p* = 0.08) were not statistically significant. The two-way interaction of day and session was also not statistically significant (*F*_1, 242_ = 2.99, *p* = 0.09). All other main effects and interactions were not significant (all *p* > 0.10).

### Relative place field characteristics for spatial information content – Spatial novelty

One-sample *t*-tests were conducted to determine if relative spatial information content between sessions differed from chance level (0.5) during the spatial novelty experiment (Fig. 8). Relative spatial information content from B to A, calculated as the ratio between the unit’s information content in B and the sum of information content in A and B, was lower than chance for both dCA1 and iCA1 on Day 1 (both *p* < 0.001), but not Day 2 (*p* > 0.10). Similarly, relative spatial information content from B to C was lower than chance for both dCA1 and iCA1 on Day 1 (*p* < 0.05 and *p* < 0.01, respectively), but not Day 2 (*p* > 0.10). Comparing session C to A, relative spatial information content was lower than chance for dCA1 on Day 1 (*p* < 0.001), but not Day 2 (*p* > 0.10). Relative spatial information content from C to A for iCA1 was not different from chance on either day (both *p* > 0.10).

**Figure 8.**
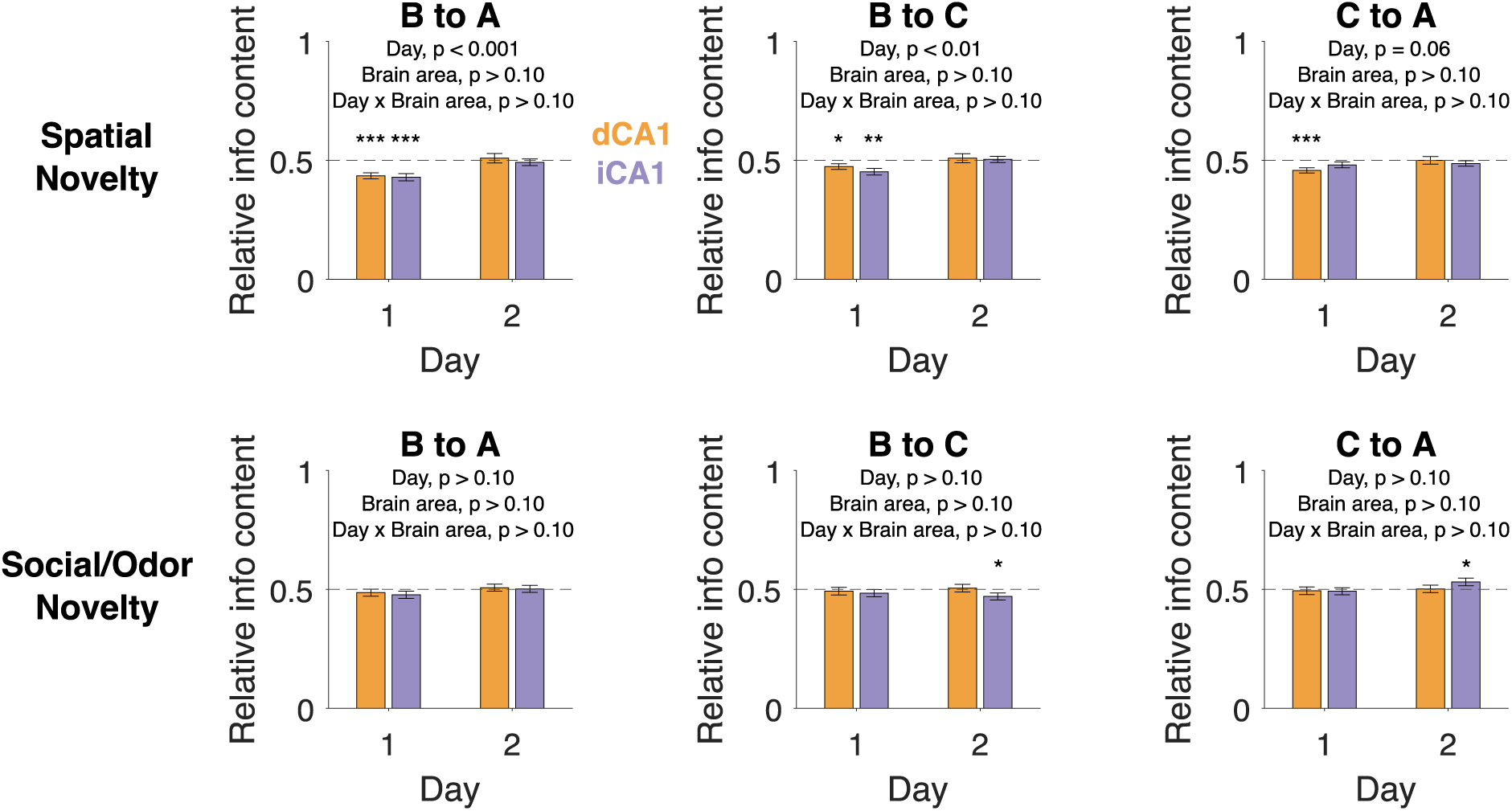
Relative information content. Relative information content for “B to A” was calculated as Session B’s information content divided by the sum of Session B and A’s information content, B/(B+A). Values below 0.5 indicate Session B’s information content is lower than chance. Relative information content calculations for B to C was calculated as B/(B+C). Relative information content calculations for C to A was calculated as C/(C+A). Error bars denote standard error of the mean.

To compare across brain regions and days we then used two-way ANOVA. Comparing sessions B to A and B to C, there was a statistically significant main effect for Day 2 having higher relative info content than Day 1 (*F*_1, 221_ = 19.18, *p* < 0.001 and *F*_1, 226_ = 8.56, *p* < 0.01). There was not a statistically significant difference across days when comparing sessions C and A (*F*_1, 221_ = 3.53, *p* = 0.06). In all three comparisons, the main effect for brain area and interaction were not statistically significant (*p* > 0.10).

### Relative place field characteristics for spatial information content – Social/odor novelty

We used a similar analysis for the novel social and odor experiments. Here we found that relative spatial information content was statistically significantly lower than chance in iCA1 when comparing B to C on Day 2 and higher than chance when comparing C to A on Day 2 (*p* < 0.05). However, in all other cases the relative spatial information content from chance did not show a statistically significant difference from chance (all *p* > 0.10), and two-way ANOVAs examining the main effects and interactions of day and brain area were not statistically significant (all *p* > 0.10).

### Relative place field characteristics for total number of place fields, field size, and active pixels

Relative place field characteristics for total number of place fields, field size, and active pixels were also calculated using the same methods above. Results for all one-sample t-tests and two-way ANOVAs are presented on Supplementary Table 1. The percentage of pixels above 20% of the peak firing rate was statistically significantly higher in iCA1 than dCA1 when comparing novel social/odor cues session B to A, but all other place field features and comparisons were not statistically significant. Overall, there were no systematic differences between dCA1 and iCA1 relative place field characteristics during novel spatial experiences or novel social/odor experiences, suggesting dCA1 and iCA1 may have similar sensitivity in processing novelty.

## Discussion

Genetic differences within the hippocampus suggest it can be divided into three regions: dHP, iHP, and vHP. The dHP receives inputs from dorsolateral entorhinal cortex and sends efferent projections to dorsal subiculum, retrosplenial cortex, dorsal lateral septum, and mammillary body, whereas, the iHP receives stronger inputs from ventromedial entorhinal cortex and sends efferent projections to prelimbic mPFC, medial olfactory nucleus, and lateral amygdala (Strange et al., 2014). This difference between the two regions could underlie possible function differences. Prior work suggested intermediate and ventral hippocampus are dispensable for spatial navigation (Moser et al., 1995), however, others have found them to be necessary (Contreras et al., 2018; Floresco et al., 1996; Loureiro et al., 2012; Wang & Cai, 2008). Importantly, iHP lesions in these studies are more likely to affect trisynaptic circuits than manipulations to the poles (Strange et al., 2014). The degree to which iHP and vHP are involved in spatial processing may also depend on experimental task demands and training protocol (Bast et al., 2009; de Hoz & Martin, 2014). Notably, in humans, there is evidence of different processing in the posterior and middle regions of the hippocampus (Dalton et al., 2018; Evensmoen et al., 2015; Poppenk et al., 2013). We based our division between dHP and iHP on a recently published single-unit study analyzing these two areas (Jin & Lee, 2021). In the current study, we analyzed global remapping between dCA1 and iCA1 neurons. Further, we measured the difference in sensitivity across different novel spatial or social/odor experiences.

### Spatial properties of dHP and iHP

Consistent with prior literature, we found differences in the raw characteristics of place field firing between dHP and iHP (Fig. 4). Place fields in iCA1 exhibited lower spatial information content, a greater total number of place fields, larger place field size, and higher percent of pixels >20% of the peak firing rate. Changes in spatial scaling may be interpreted as differences in specific, fine-grain information in dHP vs. general, coarse-grain contextual processing in more ventral regions. Nevertheless, when analyzing equal populations of units, precise spatial coding was found to be preserved in both dorsal and ventral cells (Keinath et al., 2014). Future work may seek to measure global remapping on a larger physical maze, as ventral place fields have been shown to reach ∼10 m at the ventral hippocampal pole (Kjelstrup et al., 2008). Notably, when examining spike width x firing rate x spatial information content, dCA1 and iCA1 units seemed to differ in their clustering of putative interneuron and complex spike units (Fig. 3). There seemed to be more variability in iCA1 units, whereas, dCA1 exhibited clustering similar to previous studies (Markus et al., 1994). The increased heterogeneity of cell types in the iCA1 may indicate the existence of subsets of cells unique to iCA1. Comparisons of place cell remapping between hippocampal subregions, such as dorsal CA2, suggest the area is critical for the encoding of episodic social stimuli and odor memory (Hassan et al., 2023; Oliva et al., 2020). Future investigations might aim to target the deepest region of the hippocampal ventral pole, as well as investigate further into genetic markers to demarcate subregions, cell-type diversity, and circuit level activity.

### Behavioral manipulation

The current study compared and contrasted unit activity in dCA1 and iCA1 in response to environmental novelty. To determine the strength of the novelty manipulation, we used changes in the rats’ behavior as an index. Presumably, if rats experience a large or significant event, it will interrupt their normal ongoing behavior. This would cause a change in their trial running trajectory or speed. Using this measure, we found that the spatial manipulation (novel rotation of maze arm) produced increases in the animals’ running latencies on the first day and the effect was gone on the second day, presumably because the experience was no longer novel (Fig. 5). Social/odor novelty resulted in increased exploration of the region adjacent to the novel stimulus. This effect also was gone on the second day. Similarly, we recently showed under less restrained conditions, a difference between the initial and following-day response to a familiar or novel conspecific in a spatial exploration task (Lee et al., 2022). Unlike the overt behavioral changes above, we found that even relatively modest behavioral changes were related to differences in effectiveness of observational learning (Troha et al., 2023).

### Neuronal response to the spatial manipulation

The dCA1 and iCA1 units responded similarly to spatial manipulations. This was true for changes in overall proportional firing rate (Fig. 6), direct rate map correlations (Fig. 7B), and relative spatial information content (Fig. 8), total number of fields, field size, and percent of pixels >20% of the peak firing rate (Supplemental Table 1). These data suggest that despite fundamental differences in place field characteristics (Fig. 4), both dorsal and intermediate hippocampus are similarly sensitive to spatial novelty. Associational projections within and across hippocampal hemispheres target up to two-thirds the extent of the dorsoventral axis (Amaral & Witter, 1989). Theta coherence between dHP and iHP brain areas is relatively high compared to dHP and vHP (Patel et al., 2012). When iHP was inactivated, dHP and vHP theta rhythms became uncoupled and the frequency of vHP theta decreased (Goutagny et al., 2009). Moreover, information from dHP and iHP sharp-wave ripples is integrated before being sent downstream, whereas, vHP ripples send relatively isolated information (Patel et al., 2013).

### Neuronal response to the social/odor manipulation

No remapping was found for the social/odor manipulation despite the change in the animal’s overt behavior (Fig. 5C). Interestingly, a recent paper found little evidence of remapping in dorsal or ventral hippocampus in response to social stimuli (Wu et al., 2023). Presentation of a novel male conspecific, female bedding, or coyote urine in the current study was not associated with any predictive outcomes because these occurred alongside the maze and could not be accessed by the animal. Conversely, the current study’s spatial manipulation changed the animals’ trajectory and the location of the food reward. Further, the rats were trained extensively and highly motivated to get food rewards on both ends of the linear track. Predator odors can elicit changes in behavior, though freezing might be an ineffective response to nearby predators, in which case rats might want to switch to a flight response (Eilam, 2005; M. E. Wang et al., 2012).

There is a gradient of connectivity from the hippocampus to NAc, and hippocampal units were found to code for reward locations (Gauthier & Tank, 2018). Both dHP and iHP encode spatial information but may differ in their processing of motivation and reward. Memory of specific reward locations was encoded in iHP, whereas, dCA1 encoded the memory by increasing population activity (Jarzebowski et al., 2022). The encoding of motivational significance and value changes was found to be more prominent in iHP (Jin & Lee, 2021). It is possible that assigning motivational significance to our social/odor manipulations would have resulted in remapping.

Taken together, our data support previous findings of differences in spatial firing characteristics between dorsal and intermediate cells, however, cells in both regions responded similarly to spatial and social/odor manipulations. These findings might be due to the dense redundancy and recurrent processing between the hippocampus and cortical/subcortical areas that may facilitate the integration of information. These data support a differentiation of some functions, together with an overlap of other functions along the longitudinal axis, facilitating the integration of information throughout the hippocampus (Lee et al., 2019). The current study indicated both brain areas responded similarly to novelty despite differences in their spatial tuning, suggesting they are sending out similar information but to distinct downstream targets.

## Acknowledgements

This work was supported by the University of Connecticut Research Foundation (UCRF) and the Peter and Carmen Lucia Buck Foundation (PCLB) grants to Etan J. Markus, and the Crandall-Cordero Fellowship, and The Connecticut Institute for the Brain and Cognitive Sciences Graduate Fellowship and K12 GM000680/GM/NIGMS NIH HHS/United States to Shang Lin Tommy Lee.

---

**Supplementary Table 1.**
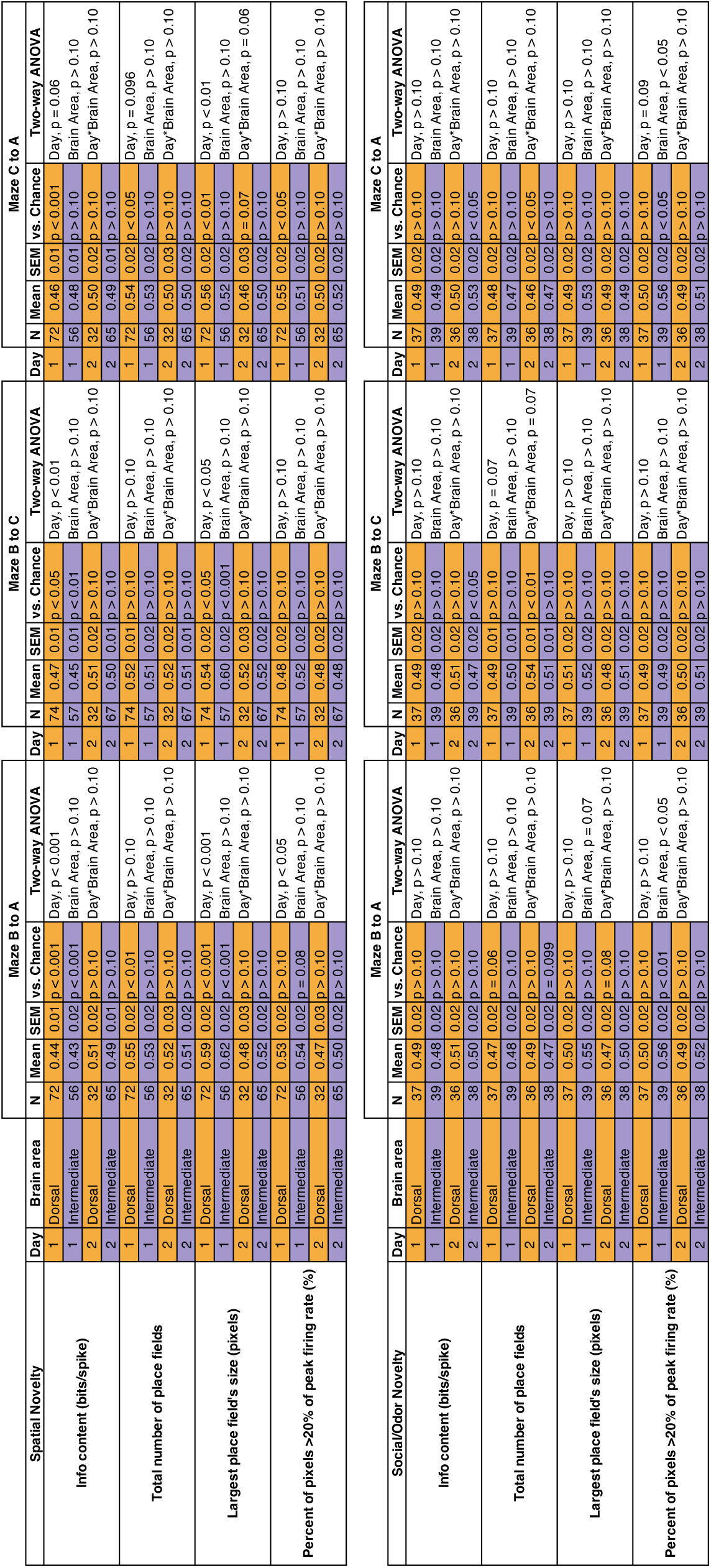
Relative place field characteristics. . The change in basic place field characteristics was calculated as the value for “Maze B to A” = Session B/(Session B + Session A), “Maze B to C” = Session B/(Session B + Session C), and “Maze C to A” = Session C/(Session C + Session A). This was done for the total number of place fields, largest place field’s size (pixels), percent of pixels above 20% of the peak firing rate, and average firing rate within the largest place field.

## References

1. Amaral, D. G., & Witter, M. P. (1989). The three-dimensional organization of the hippocampal formation: A review of anatomical data. Neuroscience, 31(3), 571–591. 10.1016/0306-4522(89)90424-7

2. Aqrabawi, A. J., & Kim, J. C. (2018). Topographic Organization of Hippocampal Inputs to the Anterior Olfactory Nucleus. Frontiers in Neuroanatomy, 12(February), 1–7. 10.3389/fnana.2018.00012

3. Bannerman, D. M., Rawlins, J. N. P., McHugh, S. B., Deacon, R. M. J., Yee, B. K., Bast, T., Zhang, W.-N., Pothuizen, H. H. J., & Feldon, J. (2004). Regional dissociations within the hippocampus — Memory and anxiety. Neuroscience & Biobehavioral Reviews, 28(3), 273–283. 10.1016/j.neubiorev.2004.03.004

4. Bast, T., Wilson, I. A., Witter, M. P., & Morris, R. G. M. (2009). From Rapid Place Learning to Behavioral Performance: A Key Role for the Intermediate Hippocampus. PLoS Biology, 7(4), 730–746. 10.1371/journal.pbio.1000089

5. Bast, T., Zhang, W.-N., & Feldon, J. (2001). The ventral hippocampus and fear conditioning in rats. Experimental Brain Research, 139(1), 39–52. 10.1007/s002210100746

6. Burwell, R. D., & Amaral, D. G. (1998). Perirhinal and postrhinal cortices of the rat: Interconnectivity and connections with the entorhinal cortex. The Journal of Comparative Neurology, 391(3), 293–321. 10.1002/(SICI)1096-9861(19980216)391:3<293::AID-CNE2>3.0.CO;2-X

7. Cenquizca, L. A., & Swanson, L. W. (2007). Spatial organization of direct hippocampal field CA1 axonal projections to the rest of the cerebral cortex. Brain Research Reviews, 56(1), 1–26. 10.1016/j.brainresrev.2007.05.002

8. Chiba, T. (2000). Collateral projection from the amygdalo-hippocampal transition area and CA1 to the hypothalamus and medial prefrontal cortex in the rat. Neuroscience Research, 38(4), 373–383. 10.1016/S0168-0102(00)00183-8

9. Chung, J. E., Magland, J. F., Barnett, A. H., Tolosa, V. M., Tooker, A. C., Lee, K. Y., Shah, K. G., Felix, S. H., Frank, L. M., & Greengard, L. F. (2017). A Fully Automated Approach to Spike Sorting. Neuron, 95(6), 1381–1394.e6. 10.1016/j.neuron.2017.08.030

10. Ciocchi, S., Passecker, J., Malagon-Vina, H., Mikus, N., & Klausberger, T. (2015). Selective information routing by ventral hippocampal CA1 projection neurons. Science, 348(6234), 560–563. 10.1126/science.aaa3245

11. Contreras, M., Pelc, T., Llofriu, M., Weitzenfeld, A., & Fellous, J.-M. (2018). The ventral hippocampus is involved in multi-goal obstacle-rich spatial navigation. Hippocampus, 28(12), 853–866. 10.1002/hipo.22993

12. Dalton, M. A., Zeidman, P., McCormick, C., & Maguire, E. A. (2018). Differentiable Processing of Objects, Associations, and Scenes within the Hippocampus. The Journal of Neuroscience, 38(38), 8146–8159. 10.1523/JNEUROSCI.0263-18.2018

13. de Hoz, L., & Martin, S. J. (2014). Double dissociation between the contributions of the septal and temporal hippocampus to spatial learning: The role of prior experience. Hippocampus, 24(8), 990–1005. 10.1002/hipo.22285

14. Dolorfo, C. L., & Amaral, D. G. (1998). Entorhinal cortex of the rat: Topographic organization of the cells of origin of the perforant path projection to the dentate gyrus. The Journal of Comparative Neurology, 398(1), 25–48. 10.1002/(SICI)1096-9861(19980817)398:1<25::AID-CNE3>3.0.CO;2-B

15. Eilam, D. (2005). Die hard: A blend of freezing and fleeing as a dynamic defense — Implications for the control of defensive behavior. Neuroscience and Biobehavioral Reviews, 29(8), 1181–1191. 10.1016/j.neubiorev.2005.03.027

16. Evensmoen, H. R., Ladstein, J., Hansen, T. I., Møller, J. A., Witter, M. P., Nadel, L., & Håberg, A. K. (2015). From details to large scale: The representation of environmental positions follows a granularity gradient along the human hippocampal and entorhinal anterior-posterior axis. Hippocampus, 25(1), 119–135. 10.1002/hipo.22357

17. Fanselow, M. S., & Dong, H.-W. (2010). Are the dorsal and ventral hippocampus functionally distinct structures? Neuron, 65(1), 7–19. 10.1016/j.neuron.2009.11.031

18. Ferguson, J. E., Boldt, C., & Redish, A. D. (2009). Creating low-impedance tetrodes by electroplating with additives. Sensors and Actuators, A: Physical, 156(2), 388–393. 10.1016/j.sna.2009.10.001

19. Floresco, S. B., Seamans, J. K., & Phillips, A. G. (1996). Differential effects of lidocaine infusions into the ventral CA1/subiculum or the nucleus accumbens on the acquisition and retention of spatial information. Behavioural Brain Research, 81, 163–171. 10.1016/S0166-4328(96)00058-7

20. Fricke, R., & Cowan, W. M. (1978). An autoradiographic study of the commissural and ipsilateral hippocampo-dentate projections in the adult rat. The Journal of Comparative Neurology, 181(2), 253–269. 10.1002/cne.901810204

21. Gauthier, J. L., & Tank, D. W. (2018). A Dedicated Population for Reward Coding in the Hippocampus. Neuron, 99(1), 179-193.e7. 10.1016/j.neuron.2018.06.008

22. Gergues, M. M., Han, K. J., Choi, H. S., Brown, B., Clausing, K. J., Turner, V. S., Vainchtein, I. D., Molofsky, A. V, & Kheirbek, M. A. (2020). Circuit and molecular architecture of a ventral hippocampal network. Nature Neuroscience. 10.1038/s41593-020-0705-8

23. Goutagny, R., Jackson, J., & Williams, S. (2009). Self-generated theta oscillations in the hippocampus. Nature Neuroscience, 12(12), 1491–1493. 10.1038/nn.2440

24. Groenewegen, H. J., der Zee, E. V.-V., te Kortschot, A., & Witter, M. P. (1987). Organization of the projections from the subiculum to the ventral striatum in the rat. A study using anterograde transport of Phaseolus vulgaris leucoagglutinin. Neuroscience, 23(1), 103–120. 10.1016/0306-4522(87)90275-2

25. Hassan, S. I., Bigler, S., & Siegelbaum, S. A. (2023). Social odor discrimination and its enhancement by associative learning in the hippocampal CA2 region. Neuron, 111(14), 2232–2246.e5. 10.1016/j.neuron.2023.04.026

26. Holt, W., & Maren, S. (1999). Muscimol Inactivation of the Dorsal Hippocampus Impairs Contextual Retrieval of Fear Memory. The Journal of Neuroscience, 19(20), 9054– 9062. 10.1523/JNEUROSCI.19-20-09054.1999

27. Jarzebowski, P., Hay, Y. A., Grewe, B. F., & Paulsen, O. (2022). Different encoding of reward location in dorsal and intermediate hippocampus. Current Biology, 1–8. 10.1016/j.cub.2021.12.024

28. Jay, T. M., & Witter, M. P. (1991). Distribution of hippocampal CA1 and subicular efferents in the prefrontal cortex of the rat studied by means of anterograde transport of Phaseolus vulgaris leucoagglutinin. Journal of Comparative Neurology, 313(4), 574–586. 10.1002/cne.903130404

29. Jimenez, J. C., Su, K., Goldberg, A. R., Luna, V. M., Biane, J. S., Ordek, G., Zhou, P., Ong, S. K., Wright, M. A., Zweifel, L., Paninski, L., Hen, R., & Kheirbek, M. A. (2018). Anxiety Cells in a Hippocampal-Hypothalamic Circuit. Neuron, 97(3), 670–683.e6. 10.1016/j.neuron.2018.01.016

30. Jin, S.-W., & Lee, I. (2021). Differential encoding of place value between the dorsal and intermediate hippocampus. Current Biology, 31(14), 3053–3072.e5. 10.1016/j.cub.2021.04.073

31. Jung, M., Wiener, S., & McNaughton, B. (1994). Comparison of spatial firing characteristics of units in dorsal and ventral hippocampus of the rat. The Journal of Neuroscience, 14(12), 7347–7356. 10.1523/JNEUROSCI.14-12-07347.1994

32. Keinath, A. T., Wang, M. E., Wann, E. G., Yuan, R. K., Dudman, J. T., & Muzzio, I. A. (2014). Precise spatial coding is preserved along the longitudinal hippocampal axis. Hippocampus, 24(12), 1533–1548. 10.1002/hipo.22333

33. Kishi, T., Tsumori, T., Yokota, S., & Yasui, Y. (2006). Topographical projection from the hippocampal formation to the amygdala: A combined anterograde and retrograde tracing study in the rat. Journal of Comparative Neurology, 496, 349–368. 10.1002/cne.20919

34. Kjelstrup, K., Solstad, T., Brun, V. H., Hafting, T., Leutgeb, S., Witter, M. P., Moser, E. I., & Moser, M.-B. (2008). Finite Scale of Spatial Representation in the Hippocampus. Science, 321(5885), 140–143. 10.1126/science.1157086

35. Komorowski, R. W., Garcia, C. G., Wilson, A., Hattori, S., Howard, M. W., & Eichenbaum, H. (2013). Ventral Hippocampal Neurons Are Shaped by Experience to Represent Behaviorally Relevant Contexts. Journal of Neuroscience, 33(18), 8079–8087. 10.1523/JNEUROSCI.5458-12.2013

36. Lee, S. L. T., Ahmed, S., Horbal, L., Pietruszewski, T., Hu, Q., & Markus, E. J. (2022). Social factors influence solo and rat dyads exploration of an unfamiliar open field. Animal Cognition, 0123456789, 3–8. 10.1007/s10071-022-01664-y

37. Lee, S. L. T., Lew, D., Wickenheisser, V., & Markus, E. J. (2019). Interdependence between dorsal and ventral hippocampus during spatial navigation. Brain and Behavior, 9(10), 1–14. 10.1002/brb3.1410

38. Lee, S. L. T., Timmerman, B., Pflomm, R., Roy, N., Kumar, M., & Markus, E. J. (2023). Sequential order spatial memory in male rats: Characteristics and impact of medial prefrontal cortex and hippocampus disruption. Neurobiology of Learning and Memory, 200(November 2022), 107739. 10.1016/j.nlm.2023.107739

39. Loureiro, M., Lecourtier, L., Engeln, M., Lopez, J., Cosquer, B., Geiger, K., Kelche, C., Cassel, J. C., & Pereira De Vasconcelos, A. (2012). The ventral hippocampus is necessary for expressing a spatial memory. Brain Structure and Function, 217(1), 93–106. 10.1007/s00429-011-0332-y

40. Maren, S., & Holt, W. G. (2004). Hippocampus and Pavlovian fear conditioning in rats: Muscimol infusions into the ventral, but not dorsal, hippocampus impair the acquisition of conditional freezing to an auditory conditional stimulus. Behavioral Neuroscience, 118(1), 97–110. 10.1037/0735-7044.118.1.97

41. Markus, E. J., Barnes, C. A., McNaughton, B. L., Gladden, V. L., & Skaggs, W. E. (1994). Spatial information content and reliability of hippocampal CA1 neurons: Effects of visual input. Hippocampus, 4(4), 410–421. 10.1002/hipo.450040404

42. Markus, E. J., Qin, Y. L., Leonard, B., Skaggs, W. E., McNaughton, B. L., & Barnes, C. A. (1995). Interactions between location and task affect the spatial and directional firing of hippocampal neurons. The Journal of Neuroscience, 15(11), 7079–7094. 10.1523/JNEUROSCI.15-11-07079.1995

43. Maurer, A. P., VanRhoads, S. R., Sutherland, G. R., Lipa, P., & McNaughton, B. L. (2005). Self-motion and the origin of differential spatial scaling along the septo-temporal axis of the hippocampus. Hippocampus, 15(7), 841–852. 10.1002/hipo.20114

44. Mikulovic, S., Restrepo, C. E., Siwani, S., Bauer, P., Pupe, S., Tort, A. B. L., Kullander, K., & Leão, R. N. (2018). Ventral hippocampal OLM cells control type 2 theta oscillations and response to predator odor. Nature Communications, 9(1). 10.1038/s41467-018-05907-w

45. Moser, E., Moser, M. B., & Andersen, P. (1993). Spatial learning impairment parallels the magnitude of dorsal hippocampal lesions, but is hardly present following ventral lesions. The Journal of Neuroscience, 13(9), 3916–3925. 10.1523/JNEUROSCI.13-09-03916.1993

46. Moser, M. B., Moser, E. I., Forrest, E., Andersen, P., & Morris, R. G. (1995). Spatial learning with a minislab in the dorsal hippocampus. Proceedings of the National Academy of Sciences, 92(21), 9697–9701. 10.1073/pnas.92.21.9697

47. O’Keefe, J., & Dostrovsky, J. (1971). The hippocampus as a spatial map. Preliminary evidence from unit activity in the freely-moving rat. Brain Research, 34, 171–175. 10.1016/0006-8993(71)90358-1

48. Oliva, A., Fernández-Ruiz, A., Leroy, F., & Siegelbaum, S. A. (2020). Hippocampal CA2 sharp-wave ripples reactivate and promote social memory. Nature, 587(7833), 264–269. 10.1038/s41586-020-2758-y

49. Patel, J., Fujisawa, S., Berényi, A., Royer, S., & Buzsáki, G. (2012). Traveling Theta Waves along the Entire Septotemporal Axis of the Hippocampus. Neuron, 75(3), 410–417. 10.1016/j.neuron.2012.07.015

50. Patel, J., Schomburg, E. W., Berényi, A., Fujisawa, S., & Buzsáki, G. (2013). Local generation and propagation of ripples along the septotemporal axis of the hippocampus. Journal of Neuroscience, 33(43), 17029–17041. 10.1523/JNEUROSCI.2036-13.2013

51. Paxinos, G., & Watson, C. (2007). The Rat Brain in Stereotaxic Coordinates Sixth Edition. Elsevier Academic Press, 170, 547–612.

52. Pentkowski, N. S., Blanchard, D. C., Lever, C., Litvin, Y., & Blanchard, R. J. (2006). Effects of lesions to the dorsal and ventral hippocampus on defensive behaviors in rats. European Journal of Neuroscience, 23(February), 2185–2196. 10.1111/j.1460-9568.2006.04754.x

53. Pitkänen, A., Pikkarainen, M., Nurminen, N., & Ylinen, A. (2000). Reciprocal connections between the amygdala and the hippocampal formation, perirhinal cortex, and postrhinal cortex in rat. A review. Annals of the New York Academy of Sciences, 911, 369–391. 10.1111/j.1749-6632.2000.tb06738.x

54. Poppenk, J., Evensmoen, H. R., Moscovitch, M., & Nadel, L. (2013). Long-axis specialization of the human hippocampus. Trends in Cognitive Sciences, 17(5), 230–240. 10.1016/j.tics.2013.03.005

55. Pothuizen, H. H. J., Zhang, W. N., Jongen-Rêlo, A. L., Feldon, J., & Yee, B. K. (2004). Dissociation of function between the dorsal and the ventral hippocampus in spatial learning abilities of the rat: A within-subject, within-task comparison of reference and working spatial memory. European Journal of Neuroscience, 19, 705–712. 10.1111/j.0953-816X.2004.03170.x

56. Poucet, B., Thinus-Blanc, C., & Muller, R. U. (1994). Place cells in the ventral hippocampus of rats. NeuroReport, 5(16), 2045–2048. 10.1097/00001756-199410270-00014

57. Rao, R. P., von Heimendahl, M., Bahr, V., & Brecht, M. (2019). Neuronal Responses to Conspecifics in the Ventral CA1. Cell Reports, 27(12), 3460–3472.e3. 10.1016/j.celrep.2019.05.081

58. Richmond, M. A., Yee, B. K., Pouzet, B., Veenman, L., Rawlins, J. N., Feldon, J., & Bannerman, D. M. (1999). Dissociating context and space within the hippocampus: Effects of complete, dorsal, and ventral excitotoxic hippocampal lesions on conditioned freezing and spatial learning. Behavioral Neuroscience, 113(6), 1189– 1203. 10.1037/0735-7044.113.6.1189

59. Risold, P. Y., & Swanson, L. W. (1996). Structural evidence for functional domains in the rat hippocampus. Science, 272(5267), 1484–1486. 10.1126/science.272.5267.1484

60. Royer, S., Sirota, A., Patel, J., & Buzsaki, G. (2010). Distinct Representations and Theta Dynamics in Dorsal and Ventral Hippocampus. Journal of Neuroscience, 30(5), 1777–1787. 10.1523/JNEUROSCI.4681-09.2010

61. Skaggs, W. E., McNaughton, B. L., Gothard, K. M., & Markus, E. J. (1993). An Information-Theoretic Approach to Deciphering the Hippocampal Code. Proceedings of the IEEE, 1990, 1030–1037. 10.1109/PROC.1977.10559

62. Strange, B. A., Witter, M. P., Lein, E. S., & Moser, E. I. (2014). Functional organization of the hippocampal longitudinal axis. Nature Reviews Neuroscience, 15(10), 655– 669. 10.1038/nrn3785

63. Suzuki, W., & Amaral, D. (1994). Topographic organization of the reciprocal connections between the monkey entorhinal cortex and the perirhinal and parahippocampal cortices. The Journal of Neuroscience, 14(3), 1856–1877. 10.1523/JNEUROSCI.14-03-01856.1994

64. Troha, R., Gowda, M., (Tommy) Lee, S. L., & Markus, E. (2023). Observational learning in rats: Interplay between demonstrator and observer behavior. Journal of Neuroscience Methods, 109807. 10.1016/j.jneumeth.2023.109807

65. Van Groen, T., & Wyss, J. M. (1990). Extrinsic projections from area CA1 of the rat hippocampus: Olfactory, cortical, subcortical, and bilateral hippocampal formation projections. The Journal of Comparative Neurology, 302(3), 515–528. 10.1002/cne.903020308

66. Wang, G. W., & Cai, J. X. (2008). Reversible disconnection of the hippocampal-prelimbic cortical circuit impairs spatial learning but not passive avoidance learning in rats. Neurobiology of Learning and Memory, 90, 365–373. 10.1016/j.nlm.2008.05.009

67. Wang, M. E., Wann, E. G., Yuan, R. K., Ramos Alvarez, M. M., Stead, S. M., & Muzzio, I. A. (2012). Long-Term Stabilization of Place Cell Remapping Produced by a Fearful Experience. Journal of Neuroscience, 32(45), 15802–15814. 10.1523/JNEUROSCI.0480-12.2012

68. Weeden, C. S. S., Roberts, J. M., Kamm, A. M., & Kesner, R. P. (2015). The role of the ventral dentate gyrus in anxiety-based behaviors. Neurobiology of Learning and Memory, 118, 143–149. 10.1016/j.nlm.2014.12.002

69. Witter, M. P., & Groenewegen, H. J. (1984). Laminar origin and septotemporal distribution of entorhinal and perirhinal projections to the hippocampus in the cat. Journal of Comparative Neurology, 224(3), 371–385. 10.1002/cne.902240305

70. Wu, W., Yiu, E., Ophir, A. G., & Smith, D. M. (2023). Effects of social context manipulation on dorsal and ventral hippocampal neuronal responses. *Hippocampus*, December 2022, 1–14. 10.1002/hipo.23507

